# Signaling in microbial communities with open boundaries

**DOI:** 10.1101/2023.01.20.524904

**Authors:** James J. Winkle, Soutick Saha, Joseph Essman, Matthew R. Bennett, William Ott, Krešimir Josić, Andrew Mugler

## Abstract

Microbial communities such as swarms or biofilms often form at the interfaces of solid substrates and open fluid flows. At the same time, in laboratory environments these communities are commonly studied using microfluidic devices with media flows and open boundaries. Extracellular signaling within these communities is therefore subject to different constraints than signaling within classic, closed-boundary systems such as developing embryos or tissues, yet is understudied by comparison. Here, we use mathematical modeling to show how advective-diffusive boundary flows and population geometry impact cell-cell signaling in monolayer microbial communities. We reveal conditions where the intercellular signaling lengthscale depends solely on the population geometry and not on diffusion or degradation, as commonly expected. We further demonstrate that diffusive coupling with the boundary flow can produce signal gradients within an isogenic population, even when there is no flow within the population. We use our theory to provide new insights into the signaling mechanisms of published experimental results, and we make several experimentally verifiable predictions. Our research highlights the importance of carefully evaluating boundary dynamics and environmental geometry when modeling microbial cell-cell signaling and informs the study of cell behaviors in both natural and synthetic systems.

SIGNIFICANCE
Microbial communities in natural environments and microfluidic devices are often exposed to open boundaries and flows, but models used to characterize diffusive signaling in such systems often ignore how device geometry, boundary conditions, and media flow influence signaling behavior. We demonstrate how the effective signaling capacity of these communities can be shaped by population geometry and advective-diffusive boundary flow in quasi-2D environments. Our approach provides a general framework to understand and control advection-reaction-diffusion systems—and their interactions with cellular signaling networks—in both natural and synthetic environments.

## INTRODUCTION

Diffusive signaling coordinates multicellular processes from embryogenesis and tissue development (1) to microbial quorum sensing (2–6). In the classical picture of diffusive signaling, the diffusible components are confined to the vicinity of the cells, either by external barriers such as an embryonic envelope, or by the cell membranes themselves in the case of direct cell-to-cell molecular exchange. In these cases, global signaling properties are determined by basic transport parameters such as the diffusivity of the signaling molecule or the speed of advective flow within the cell population (7).

However, in other cases, the diffusible components are free to escape at the population boundaries, or are otherwise affected by properties of the surrounding medium such as fluid flow. These cases include microbial communities such as biofilms or swarms, whose boundaries are usually open and dynamic, and which often form at the interfaces of solid substrates and fluid flows (8). In such environments, responses at the macroscopic (population) level depend on both the features of the domain in which the cells grow and the dynamics of the constituent cells. In particular, open boundaries can significantly impact signaling behavior within the community. Such impacts include modulation of signaling depth and spatial signaling profiles (9), as well as the challenges that signaling systems face when trying to respond to time-varying flows in a spatiotemporally robust manner (8, 10). Despite the importance of these impacts, signaling in open geometries has been understudied relative to signaling in closed geometries. As we will show, open geometries can induce counterintuitive signaling characteristics.

Laboratory experiments aimed at characterizing signaling in microbial communities often rely on microfluidic devices. Such experiments allow researchers to characterize the behavior of spatially extended systems, thereby facilitating the design of microbial consortia that maintain desired population fractions (11) or produce emergent spatiotemporal patterns (12). Here signaling provides the necessary intercellular communication pathway to coordinate responses and achieve population-level phenotypes. In typical microfluidic experiments, cells are forced to grow in a monolayer, thereby allowing for imaging of large populations at high resolution. Such imaging capability facilitates the investigation of consortia-scale spatiotemporal dynamics, emergent collective behavior, and nematic effects (13, 14). Importantly, many microfluidic devices employ open boundaries between the cell population and the surrounding fluid in order to supply media to cells and remove waste products and excess cells. Open boundary geometries can strongly impact the dynamics of growing microbial collectives and therefore place such microfluidic devices into the same understudied paradigm as the aforementioned biofilms and swarms.

Here, we use mathematical modeling to investigate the effects of open, advective boundaries on cell-cell signaling within a bacterial monolayer. Surprisingly, in contrast to the closed-boundary case, we find that the spatial extent of signaling from a source cell does not depend on the diffusion coefficient, but rather depends entirely on the population geometry. When the signal can degrade, we find that the signaling extent is determined by the minimum of the geometric length-scale and the classical lengthscale set by the ratio of diffusion to degradation. Further, we find that flow at the boundary can introduce signal gradients within the population—even if flow is absent within the population itself—due to the diffusive exchange of signaling molecules with the boundary region. We compare our results to published data on bacterial monolayers in a microfluidic device that signal via a quorum-sensing factor.

## RESULTS

We consider a two-dimensional continuum model of bacteria in a monolayer (Fig. 1A). Such monolayer configurations are typical of the leading edge of growing colonies or biofilms (15–17) and are often imposed by thin cell-trapping regions in microfluidic devices (4, 5, 9) (Fig. 1B). To investigate boundary effects, we confine the monolayer to a rectangle with width *w* and height *L*. These parameters can be thought of as the characteristic lengthscales of a natural population, or as the precise dimensions of the rectangular trapping region in a microfluidic device. We specify the boundary conditions in detail in the following sections.

**Figure 1:**
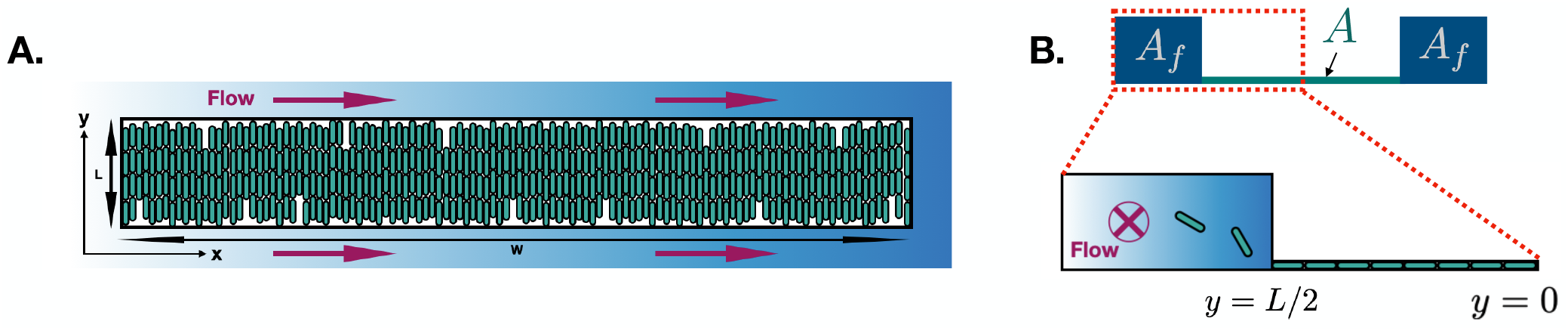
We model bacteria growing in a spatially extended microfluidic device using a continuum framework. (A) Top view of the device. Cells are confined to a thin trapping region 𝒯 of width *w* and length *L*, wherein they grow in a monolayer. The trapping region interfaces with two three-dimensional flow channels, an upper flow channel ℱ_+_ and a lower flow channel. ℱ_−_ The boundaries between the trapping region and the flow channels are open and therefore subject to flow effects. (B) Side view of the device. The trapping region has cross-sectional area *A*. Flow channels each have cross-sectional area *A* _*f*_.

To investigate intercellular signaling, we suppose that cells secrete a molecule at a rate *α* that can diffuse with coefficient *D* and degrade with rate *γ*. Secretion of diffusible molecules is a ubiquitous signaling strategy in bacteria, employed in natural functions such as quorum sensing (18, 19), and probed or engineered in microfluidic experiments (4, 5, 20). In the following sections, we investigate the effects of open boundaries and fluid flow on the properties of such signaling in steady state.

### Open boundaries make the signaling lengthscale independent of diffusion

We first investigate the spatial extent of signaling from a given source cell, without flow; we will consider flow in a subsequent section. To investigate the spatial extent of signaling, it is often convenient to consider some cells as “senders” of the signaling molecules and other cells as “receivers.” For example, in a system with closed boundaries, such as a developing embryo, signaling molecules are “sent” from one end of the embryo and “received” by cell nuclei in the bulk of the embryo (7). Similarly, in microbial communities, often one subpopulation of cells secretes a diffusible signal that a second subpopulation receives, as has been realized and studied in microfluidic experiments (9, 21).

In the case of a developing embryo, the classic synthesis-diffusion-clearance model (7) predicts that the signal will establish an exponential concentration profile in steady state. With molecular diffusion coefficient *D* and degradation rate *γ*, this profile will have a lengthscale, 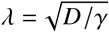, given that the system is wide (*w* ≫ *λ*). However, with open boundaries, molecules can leave the system not just by degrading but also by diffusing across the boundary. It is thus unclear what effect open boundaries will have on the signaling lengthscale.

To address this question, we consider the sender-receiver system shown in Fig. 2A, where cells in the left half of the trapping region 𝒯 (*x <* 0) produce the diffusible signal, and cells in the right half (*x*≥ 0) do not. The signal concentration *c* (*x, y*) therefore obeys the diffusion equation with production rate *α*,

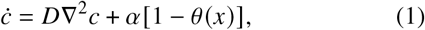

where the Heaviside function *θ* (*x*) ensures that signal production only occurs for *x <* 0. Because we are interested in open boundaries, where molecules can be lost by diffusion, degradation is not required for *c* to reach steady state. Signaling molecules often have a large half-life, and are not absorbed into microfluidic device material (22). Therefore, we neglect degradation here, and we consider its effects in the next section.

**Figure 2:**
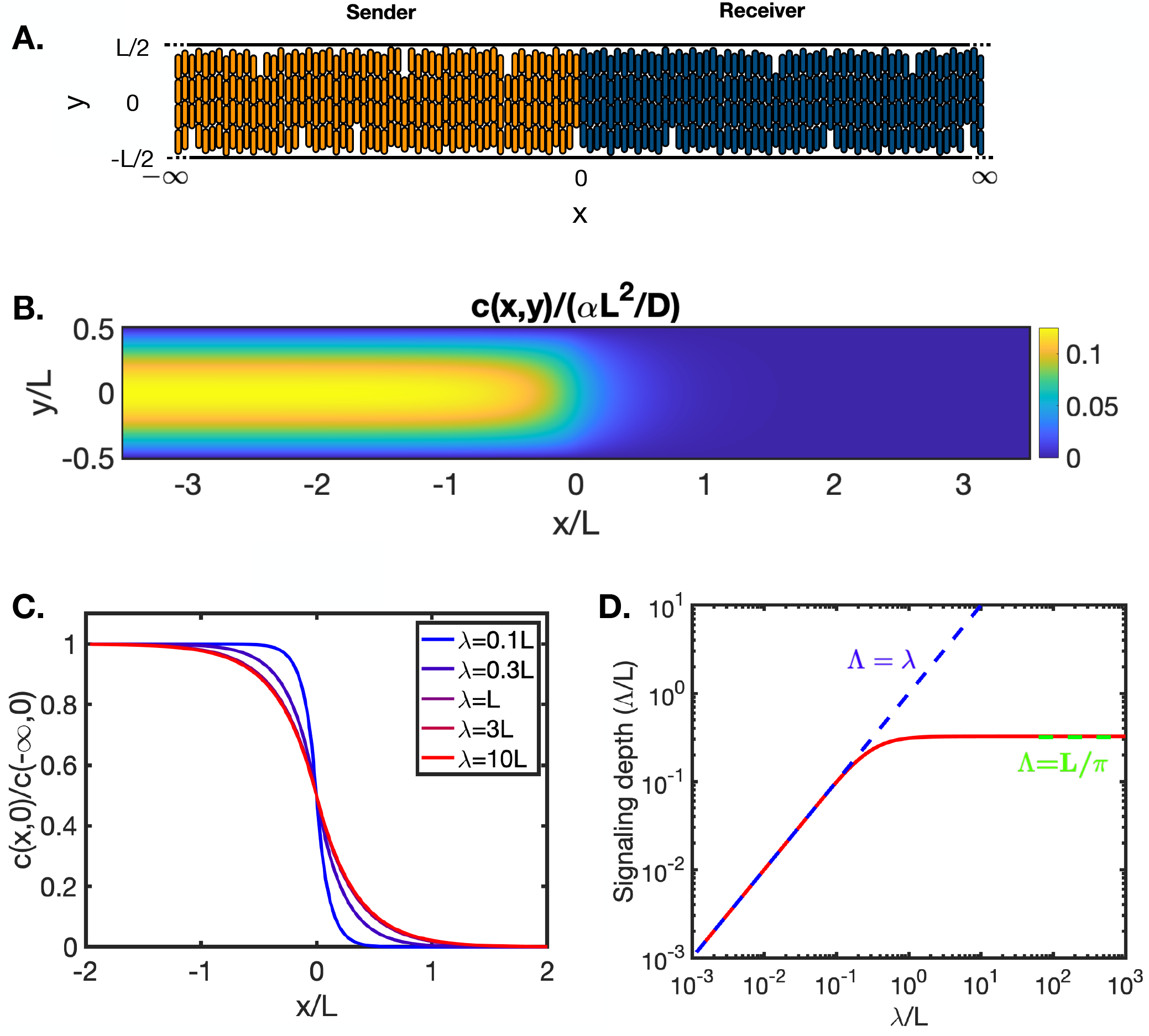
Signaling lengthscale in a “sender-receiver” setup. (A) Cells on the left (orange) secrete a diffusible signal, whereas cells on the right (blue) do not. (B) Steady-state concentration profile with no signal degradation (Eq. (5)). (C) Signal profile at the midline (*y* = 0) with degradation rate *γ* (Eq. (9)), characterized by diffusion lengthscale 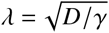. (D) Signaling is limited by the minimum of *λ* and *k L*/*π*, where *L* is the lengthscale of the trapping region (see A) and *k* ≈ 1.

Eq. (1) does not capture diffusion of signaling molecules through the cell membrane, but rather describes the dynamics of signaling molecule concentration in extracellular space. A more detailed model would include separate PDEs for intracellular and extracellular compartments coupled via diffusion through the cell membrane. However, here we capture the combined effects of molecule production and secretion into the extracellular space with a single rate, *α*. Eq. (1) then describes diffusion of signaling molecules within the extracellular space.

Eq. (1) also neglects cell growth. We are interested in the diffusion of molecules over distances *L* on the order of a hundred microns, with diffusion coefficient *D* on the order of hundreds of microns squared per second. We therefore neglect cell growth because the typical diffusion timescale *L*^2^ /*D* is tens of seconds, which is much smaller than the typical growth and division timescale of tens of minutes to hours.

To determine the effect of open boundaries, we assume that the upper and lower boundaries (*y* = ±*L/* 2) between the trapping region and the flow channels are absorbing with respect to the signaling molecule, so that the signal concentration vanishes there:

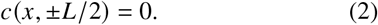

We will see later that absorbing boundary conditions are appropriate if we assume the flow channels have large cross-sectional dimensions, an assumption we make for our sender-receiver analysis. Since we are modeling a wide system (*w*≫ *L*), we take *w* → ∞. A sender-receiver system of precisely this type (with open boundaries at *y* = ± *L/* 2 and with *w≫ L*) was created in microfluidic experiments in previous work (9). We compare our predictions to these experiments in the Discussion.

Far from the sender cells, the signal concentration must vanish:

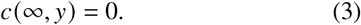

Far from the receiver cells, the concentration must become independent of *x*. The diffusion equation there reads 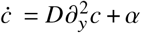, which in steady state and with Eq. (2) is solved by

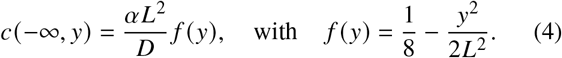

With the four boundary conditions in Eqs. (2)–(4), we solve Eq. (1) in steady state by separating variables and ensuring continuity of the solution and its derivative at the sender-receiver interface (Appendix A). The result is

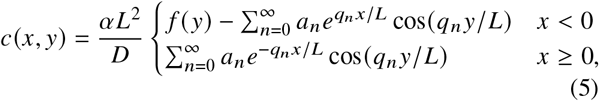

where 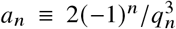 and *q*_*n*_ ≡ *π*(2*n* + 1). The signal concentration given by Eq. (5) is shown in Fig. 2B. The concentration vanishes at the open boundaries and decays across the sender-receiver interface.

The signaling lengthscale is set by the decay length of the concentration profile in the right half (*x*≥ 0) of the trapping region. The decay length is obtained by integrating the profile, normalized by its value at the interface (*x* = 0). Doing so along the midline (*y* = 0), we obtain a measure of signaling depth,

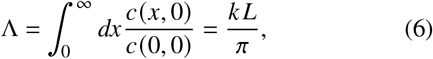

where 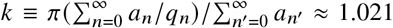. Eq. (6) thus shows that the signaling lengthscale Λ depends only on *L*, the height of the trapping region between the two open boundaries. Surprisingly, it is independent of the diffusion coefficient, *D*. In fact, the lack of a diffusion-dependent lengthscale is already apparent in Eq. (5), where we see that *D* factors out of the solution and is absent from the exponential *x*-dependence. Thus the diffusion coefficient affects the overall amplitude of the signal, but not how it decays with distance from the sender-receiver interface.

The previous observations elucidate the mechanism behind the diffusion-independence of the signaling depth. When diffusion increases, the amplitude of the profile decreases, in both open and closed systems, as diffusion spreads the signaling molecules across space. In closed systems, this spreading leads to an increased signaling lengthscale as higher diffusion allows molecules to travel farther from the source. However, in open systems, increasing the spread of molecules across space also increases the rate at which they cross the open boundaries. Molecules that would have diffused farther from the source if boundaries were closed are the ones that are more likely to be lost. In systems with open boundaries, the two opposing effects of signal loss across the open boundaries and signal diffusion cancel exactly, so that the signaling lengthscale is independent of diffusion.

### With degradation, the signaling lengthscale is bounded from above by the geometry

Degrading enzymes can introduce active degradation in microbial populations. For example, expression of AiiA lactonase can significantly accelerate the degradation of signaling molecules inside bacteria (23), which can translate to an effective extracellular molecule loss for sufficiently fast diffusion across the membrane. Moreover, extracellular signal degradation can be induced and controlled in engineered microbial consortia (6, 24).

To examine how degradation impacts signaling in open geometries and to more directly compare the behavior of our model to that of the closed-boundary synthesis-diffusion-clearance model, we introduce degradation with rate *γ* into Eq. (1):

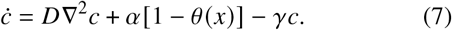

The solution far from the receiver cells generalizes from Eq. (4) to

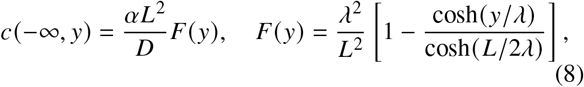

where 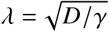 is the closed-boundary diffusion lengthscale. We use *c* (−∞, *y*) as a boundary condition for Eq. (7), together with the boundary conditions given in Eqs. (2)–(3). Solving Eq. (7) using the same approach as in the previous section (Appendix A), we obtain

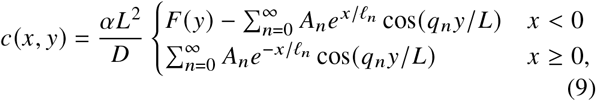

where *A*_*n*_ = 2 (−1 _*n*_ (*ℓ*_*n*_ */L*)^2^/*q*_*n*_ and 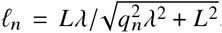. Note that in the zero-degradation limit (*λ ≫L*), we have *F* (*y*) → *f* (*y*), *ℓ*_*n*_→ *L /q*_*n*_, *A*_*n*_ →*a*_*n*_, and Eq. (9) reduces to Eq. (5), as expected.

The solution given by Eq. (9) is shown in Fig. 2C, where we see that the signal penetrates more deeply into the receiver-cell population as *λ* increases, but that the signaling depth eventually saturates. Indeed, with degradation the penetration depth of the signal defined in Eq. (6) becomes

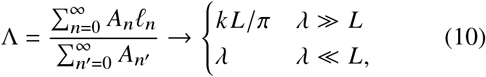

where the second case follows from the fact that *ℓ*_*n*_ → *λ* for *λ ≫L*. Eq. (10) is plotted in Fig. 2D. We recover the diffusion-independent result given by Eq. (6) in the limit of small degradation, *λ* ≫ *L*. In this limit, signal loss due to diffusion into the flow channels dominates over signal loss due to degradation. When degradation is strong, *λ ≪L*, signaling depth approaches the closed-boundary lengthscale, *λ*. In this limit, signaling molecules typically degrade before they diffuse into the flow channels. Overall, we have

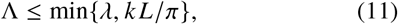

a bound that reflects the influence of both degradation and open boundaries on signaling depth.

### Flow outside the population introduces signal gradients within the population

Thus far we have considered the effects of open boundaries, but not fluid flow in the boundary regions. Surrounding flows are common in natural settings (8, 10), and flow is often desired or operationally necessary in channels bounding the trapping region in a microfluidic device (25, 26). To investigate the effects of boundary flow on signaling in a bacterial population, we return to the simplest case of a homogeneous population (all cells secrete the signal) with no signal degradation (Fig. 3A). We introduce flow at a constant velocity *v* in the *x*-direction within the flow channels ℱ_±_ that lie outside the upper and lower boundaries of the trapping region (*y* = ±*L/* 2). Because flow breaks the translational symmetry of the signal profile in the *x*-direction, we assume the width of the trapping region, *w*, is finite. Specifically, we allow the trapping region to extend from *x* = 0 to *x* = *w* and impose reflective conditions at these boundaries:

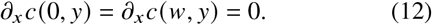

We will see later that the signal profile in the flow channels (and thus the trapping region) is largely insensitive to the boundary conditions at these ends.

**Figure 3:**
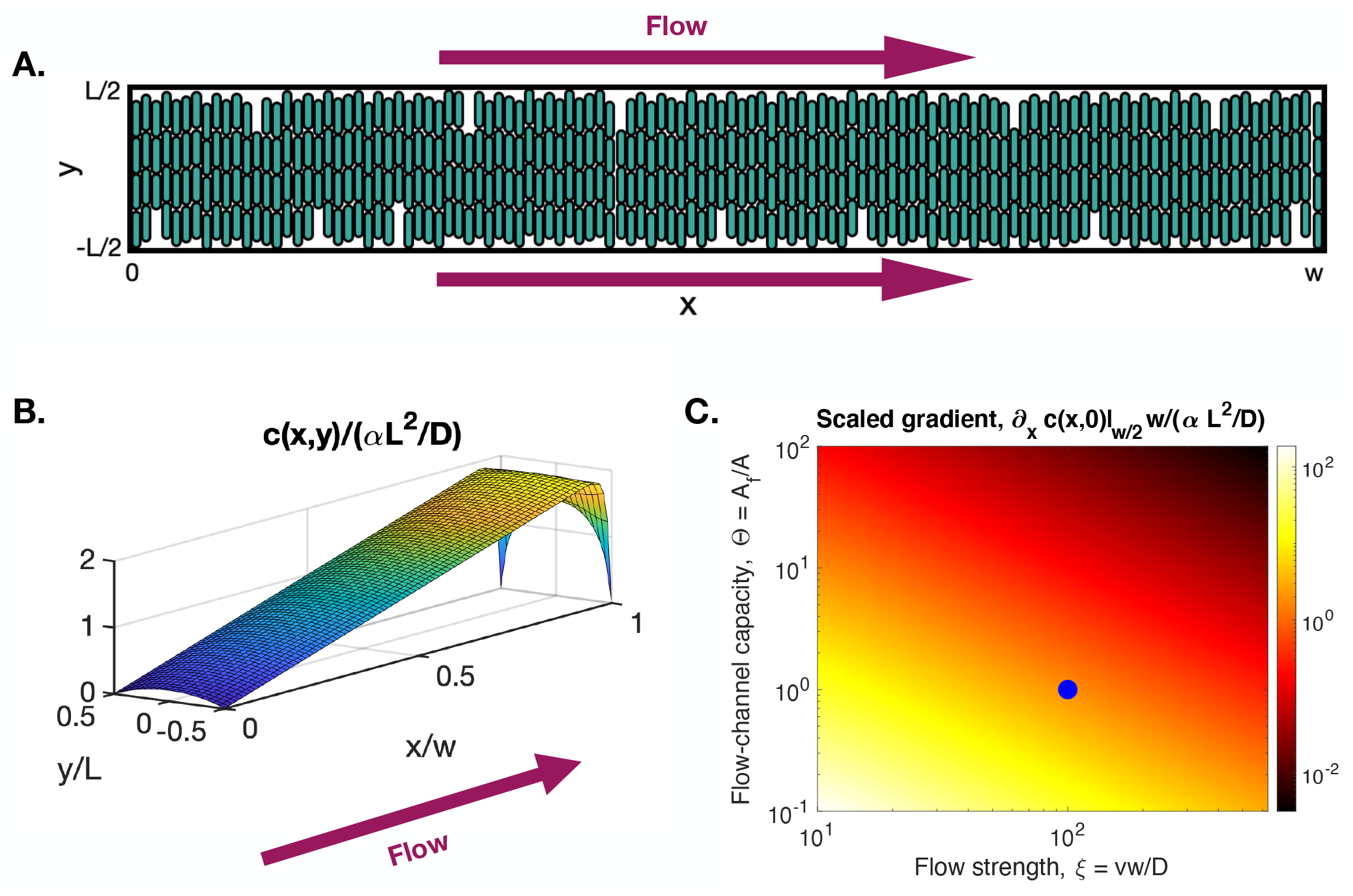
Effects of boundary flow. (A) Rightward flow is imposed at the upper and lower boundaries. (B) The flow induces a concentration gradient not just at the boundaries (long edge), but also within the cell population (surface). (C) The gradient is largest for intermediate flow strength *ξ* and small flow-channel capacity Θ. B and C are plots of Eq. (19). Parameters are *ξ* = 100, Θ = 1, and aspect ratio *ϕ* = *w*/*L* = 20 in B; and *ϕ* = 20 in C. Plot in B corresponds to blue circle in C.

Since we model the trapping region 𝒯 as a two-dimensional domain, it is convenient to average over the *z*-direction in the flow channels. Indeed, this type of dimension reduction is often performed when studying pollutant transport in rivers (27) or shallow-water flows (28). Let *b*_±_ (*x, y*) denote the concentration of signaling molecule in ℱ_±_, averaged over the *z*-direction. The dynamics in the trapping region and flow channels obey

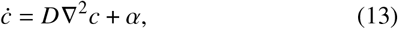

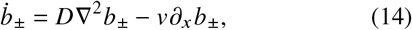

where *α* is the signal production rate and *v* is the flow velocity. Although cells can exit the trapping region and enter the flow channels, we assume that signal production in the channels is negligible. We also assume that there is no flow in the trapping region. However, the trapping region and the flow channels are coupled by diffusion of molecules across the boundaries. Correspondingly, we impose continuity of the profiles at the boundaries,

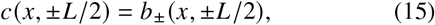

as well as their derivatives, *∂*_*y*_ *c*|_*y*=±*L/*2_ = *∂*_*y*_ *b*_±_|_*y*=± *L*/2_.

To solve Eqs. (13) and (14), we assume that whereas the concentration in the flow channels is heterogeneous in the *x*-direction due to the flow, it is homogeneous in the *y*-direction, so that *b*_+_ (*x, y*) = *b*_−_(*x, y*) = *b* (*x*). Such an approximation is valid when the length of the flow channels is an order of magnitude larger than their width (27). Because *b* no longer depends on *y*, the net flux of signal into the flow channels can no longer be accounted for by enforcing continuity of the *y*-derivative at the boundaries. Instead, this flux appears as an effective source term in Eq. (14) whose magnitude is determined by flux balance (the validity of this argument will be addressed *post hoc* at the end of this section). Specifically, in a slice of width Δ*x*, the flux of signaling molecules out of the trapping region, *α A*Δ*x*, must equal the flux into the two flow channels, 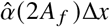, where *A* and *A* _*f*_ are the cross-sectional areas of the trapping region and each flow channel, respectively (Fig. 1B). Thus, the effective source term is 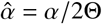 for the area ratio Θ = *A*_*f*_/*A*, which we refer to as the flow-channel capacity. Correspondingly, Eq. (14) becomes

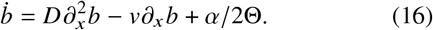

We impose absorbing boundary conditions on the flow channels at either end,

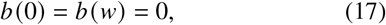

which corresponds to rapid removal of signaling molecules there. In Appendix B, we show that for long flow channels with sufficiently fast flow, the profile *b*(*x*) in the bulk is insensitive to the boundary conditions.

We solve Eqs. (13) and (16), with the boundary conditions in Eqs. (12), (15), and (17), using separation of variables (Appendix C). The result is

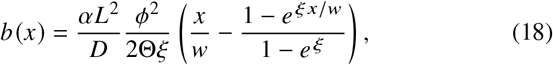

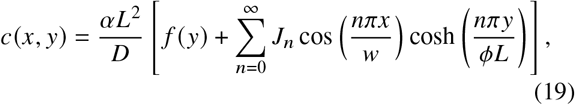

where

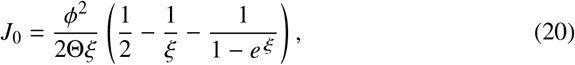

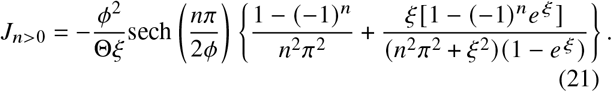

Here, *f* (*y*) is as in Eq. (4), *ξ* = *vw*/*D* is the Péclet number of the flow channel (a dimensionless determinant of the flow strength relative to diffusion), and *ϕ* = *w*/*L* is the aspect ratio of the trapping region. Note that for either *Θ* → ∞ or *ξ* → ∞, Eq. (18) reduces to *b* (*x*) = 0, and Eq. (19) reduces to Eq. (4) because *J*_*n*_→ 0. The reduction occurs because in either of these limits—very large flow channels or very fast flow, respectively—molecules leaving the trapping region never return, and the flow channel becomes an absorbing boundary.

Eqs. (18) and (19) are shown, for representative values of *ξ, Θ*, and *ϕ*, in Fig. 3B: *c (x, y)* is the surface and *b (x)* is the long edge. We see that the concentration increases in the flow channels along the flow direction. For *ξ* ≫ 1, the increase in *b(x)* is linear in *x*, sufficiently far from the boundaries at *x* = 0 and *x* = *w* (Appendix B). We see that the concentration increases not only in the flow channels (the edge), but also within the cell population (the surface). Thus, diffusive coupling between the flow channels and the trapping region induces a signal gradient in cell population, even though the population itself is not subjected to the flow.

To get a sense of the magnitude of the gradient within the cell population, we plot in Fig. 3C the derivative ∂ _*x*_ *c* (*x, y*), scaled by the characteristic lengthscale *w* and concentration value *αL*^2^/*D*, and evaluated at the midpoint of the trapping region, *x* = *w*/2 and *y* = 0, as a function of the flow strength *ξ* and the flow-channel capacity 0. We see that the gradient vanishes in the two absorbing-boundary limits mentioned above (Θ→ ∞ and *ξ*→ ∞). On the other hand, the gradient can be large for flow of intermediate strength and channels of limited capacity. For example, the case plotted in Fig. 3B, corresponding to the blue circle in Fig. 3C, has parameters estimated from recent microfluidic experiments with *E. coli* (9), and we see that the gradient is substantial. We comment further on this point in the Discussion.

Our solution Eq. (19) relies on the validity of the effective source term *α*/ 2Θ in Eq. (16). This term is a local approximation in *x* for the rate of increase of flow channel concentration due to diffusive coupling with the trapping region under a flux balance argument. Our use of the effective source term is therefore an approximation, which we validate by computing the transverse flux, −*D* ×∂_*y*_ *c* × *δΔx*, evaluated at the boundary *y* = ±*L*/2, where the trap depth is *δ* = *A*/*L*. Indeed, considering only the term *f* (*y*) in Eq. (19), we have

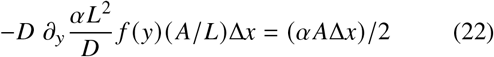

at this boundary location, which after volume scaling is equivalent to the effective source term *α*/2Θ. Thus, the additional flux due to the series solution in Eq. (19) is a residual flux that violates our original assumption regarding the validity of the effective source term.

To simplify our model, enforce the flux boundary coupling, and obtain a prediction that the concentration gradient within the cell population is linear in *x* in the bulk (away from the left and right boundaries), we replace the series solution in Eq. (19) with the linear flow channel approximation

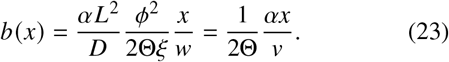

(This is Eq. (43) from Appendix B.) This enforces the flux balance approximation since the diffusion operator in Eq. (13) satisfies ∇^2^ (*c*(*x, y*) + *b* (*x*)) = ∇^2^*c*(*x, y*) when *b* is linear. The resulting *c*(*x, y*) is quadratic in *y* and linear in *x*:

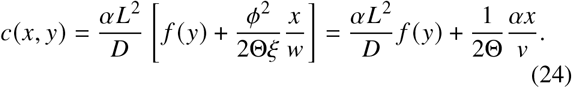

Here *f* (*y*) is given by Eq. (4). Note that this simplification is self-consistent in that it satisfies our flux boundary coupling assumption.

Eq. (24) suggests that in the bulk (away from the left and right boundaries), boundary flow induces a linear (in *x*) gradient within the cell population. Fig. 3B is consistent with this prediction.

## DISCUSSION

We described and analyzed a tractable model to explain how advective-diffusive boundary conditions shape signaling response in spatially-extended microbial communities. We assumed bacteria are trapped in a monolayer within a region bounded by two adjacent channels through which fluid flows. In the limit of zero flow speed with large, absorbing channels, we found that the signaling lengthscale is determined (or, with degradation, bounded) by the monolayer geometry, not the diffusion coefficient, because diffusion disperses molecules but also hastens their loss at the boundaries. We also found that flow at the boundaries can induce significant signal gradients in the population and that this effect is most pronounced with small flow channels at intermediate flow speeds. Although we based the model on a microfluidic trap setting, a similar approach can be used to describe more general situations. For instance, a thin bacterial film growing in a pipe could be modeled by assuming that an adjacent channel lies above a layer of cells.

Our results could have significant impact on quorum sensing in microbial populations. A principal function of quorum-sensing (QS) circuits in natural systems is the detection of a quorum of cells that triggers induction of a gene network. For example, a QS signal can trigger the production of proteins that release the extra-cellular matrix so that cells move to a mobile state under starvation (2). Pai and You have described this as the QS circuit’s sensing potential, which depends on the local environment and a threshold level of signaling molecule sensed by the cell (3). Our results can be used to generalize this sensing potential framework to include environmental influence on QS activation. We did not model cellular responses to the QS signal, but assumed that cells that express the signal do so uniformly. Bacteria can respond to QS signals in complex ways, however. Dalwadi and Pearce have used a model similar to the one we analyzed to show that positive feedback can act as a low-pass filter and ensure a robust collective response to oscillatory flow (8). In their model the flow passes over the surface of a cell population trapped in a pocket. Their analytical results are based on the assumption that diffusion across this surface dominates the diffusion in the direction of the flow, allowing them to derive a tractable one-dimensional PDE for the signal concentration in the direction perpendicular to the flow.

Our model provides several experimentally testable predictions. First, for bacterial collectives growing in geometries with open boundaries, chemical signaling depth can be independent of the diffusion rate of the signaling molecule. Second, when a flow channel borders a bacterial collective, signaling molecule flux into the flow channel can induce a graded signal concentration profile there. This graded profile in the flow channel can induce signaling molecule concentration gradients *within the bacterial collective*, even when the bacterial collective is isogenic. In this way, flow may play a role in differentiation. These predictions are testable, as bacteria such as *E. coli* can be engineered to respond to the presence of a quorum-sensing signal by producing a fluorescent protein in a graded manner, or when signal concentration reaches a threshold.

As a first step in comparing our results to experiments, we can consider a previous study in which a sender-receiver system of the type in Fig. 2A was constructed in a microfluidic device (9). The height of the trapping region was *L* = 100 *μ*m, from which Eq. (6) predicts that the signal should extend for a lengthscale of Λ ≈*L/π* ≈32 *μ*m. The measured lengthscale was Λ = 20 *μ*m, which agrees within a factor of two. The prediction could be refined by considering the effect of boundary flow (Fig. 3) on the sender-receiver geometry (Fig. 2), which could conceivably increase the predicted lengthscale (the experimental parameters *w* = 2000 *μ*m, *D* = 500 *μ*m^2^/s, *v*∼ 25 *μ*m/s, *A* = 100 *μ*m ×1 *μ*m, and *A* _*f*_ = 10 *μ*m ×10 *μ*m give *ξ* = *vw* /*D* = 100, Θ = *A*_*f*_ /*A* = 1, and *ϕ* = *w*/ *L* = 20, as in Fig. 3B). On the other hand, the fact that cells in the experiment are nematically ordered with their long axis pointing toward the open boundaries, as in Fig. 2A (13, 14), could conceivably decrease the predicted lengthscale because diffusing molecules are subject to steric barriers more often in the *x*-direction than in the *y*-direction. Even without these refinements, it is encouraging that our prediction is close to the experimental observation.

Our modeling could be extended, for example to include diffusion of signaling molecules across the cell membranes. Currently we assumed that cell-internal and cell-external signaling molecule concentrations are equal at steady state. This is tantamount to assuming that the diffusion rate of signaling molecules through the cell membrane, *d*, is infinite. When *d* is low, however, cross-membrane timescale, which scales as *d*^−1^, can become important. First, when 0 *< d <*∞, in steady state cell-internal and cell-external signaling molecule concentrations will differ in the trapping domain. This difference will increase as *d* decreases. Second, when the cell membrane is impermeable (*d* = 0), cells will sequester all of the signaling molecules they produce before said cells exit the trapping region, resulting in no signaling through the extracellular space. This sequestration effect will continue to limit cell-cell signaling efficacy when *d >* 0, provided the *d*^−1^ timescale is long relative to other system timescales. We anticipate that this and other modeling advances can be included in future work.

## APPENDIX A

Here we derive Eq. (9). As mentioned in the main text, Eq. (5) follows in the limit of no degradation (*λ ≫L*).

In the receiver domain (*x* ≥0), the source term *α* is absent from Eq. (7). Separating variables as *c* (*x, y*) = *X* (*x*) *Y* (*y*), Eq. (7) in steady state in this domain becomes

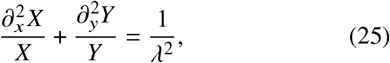

where 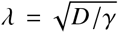. Because the two terms on the left are each a function of a different variable, and both sum to a constant, they must each equal a constant themselves. Calling the second term’s constant −*q*^2^/*L*^2^ for some unknown *q*, we have 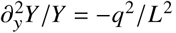, whose general solution is

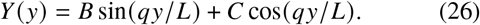

The boundary conditions in Eq. (2) require that *B* = 0 and *q* = *π* (2*n* + 1) ≡ *q*_*n*_ for nonnegative integer *n*. The first term in Eq. (25) then satisfies 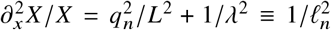, whose general solution is

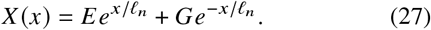

The boundary condition in Eq. (3) requires that *E* = 0. Thus, the solution in the receiver domain is

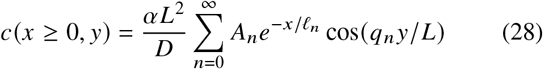

for some unknown *A*_*n*_.

In the sender domain (*x <* 0), the source term *α* is present in Eq. (7). We write the steady state solution in this domain as the sum of a particular solution, which is any function that satisfies Eq. (7) with *α* present, and the homogeneous solution, which satisfies Eq. (7) with *α* absent. For the particular solution we use the limit far from the receiver cells, Eq. (8). For the homogeneous solution, we use Eqs. (26) and (27), where again *B* = 0, but in this domain *G* = 0 to prevent the solution from diverging as *x* → −∞. Thus,

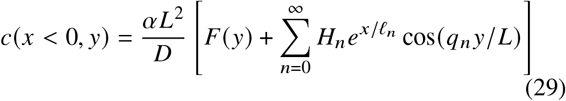

for some unknown *H*_*n*_, where *F(y)* is as in Eq. (8).

Differentiating Eqs. (28) and (29) with respect to *x*, we see that continuity of the derivative at *x* = 0 requires that *H*_*n*_ = −*A*_*n*_ for all *n* due to the orthogonality of the cosines. Continuity of the solution at *x* = 0 then requires

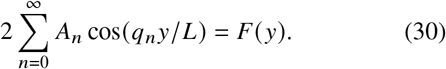

The orthogonality of cosines, expressed as

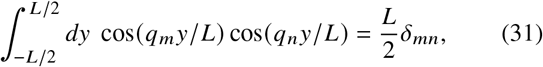

allows us to invert Eq. (30),

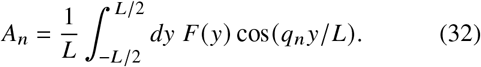

Inserting *F* (*y*) from Eq. (8) and evaluating the integrals yields

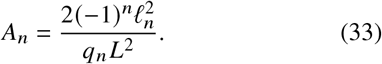

This completes the derivation of Eq. (9).

## APPENDIX B

Here we show that, for long channels with sufficiently fast flow, the profile *b* (*x*) in the bulk is linear and insensitive to the boundary conditions. We will illustrate this point by using two different choices for the boundary condition at *x* = *w* and showing that the bulk profile is the same linear function of *x* for both choices.

The bulk is defined by values of *x* that are small compared to the size of the system in that direction, *w*, but large compared to the characteristic lengthscale of the system. The characteristic lengthscale is a function of diffusion and flow speed and consequently required by dimensional analysis to scale as *D /v*. Thus, the bulk is defined by *D/ v≪ x ≪w*. Dividing by *w* and recalling that *ξ* = *vw* /*D*, this expression becomes

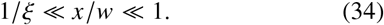

Defining *ϵ* = *x*/*w*, the two conditions in Eq. (34) become

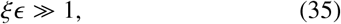

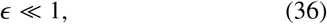

respectively. For a bulk regime to exist, the two extremes in Eq. (34) must be well separated, and therefore we must also have 1/*ξ* ≪1, or

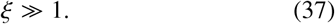

Given that *ξ* = *vw /D*, Eq. (37) makes clear that a bulk regime exists for sufficiently long channels (large *w*) with sufficiently fast flow (large *v*).

In terms of *ϵ*, Eq. (18) reads

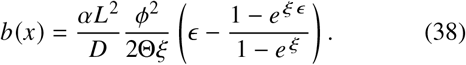

Eqs. (35) and (37) allow us to neglect the ones in the numerator and denominator, respectively, giving

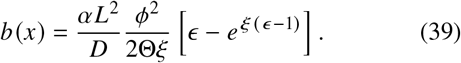

Eq. (36) allows us to neglect the *ϵ* in the exponent, and Eq. (37) then allows us to neglect the exponential altogether, giving

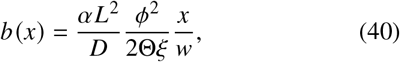

where we have restored *ϵ* = *x/ w*. We see that *b (x)* is a linear function of *x* in the bulk.

Eq. (18) satisfies *b w* = 0, but alternatively we may have the no-flux boundary condition ∂_*x*_ *b*|_*x*=*w*_ = 0, corresponding, for example, to a concentration reservoir beyond *x* = *w*. The solution of Eq. (16) that satisfies this condition, along with *b* (0) = 0, is

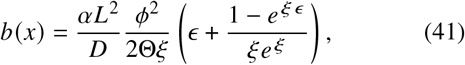

in terms of *ϵ* = *x*/ *w*. Again, Eq. (35) allows us to neglect the one in the numerator, giving

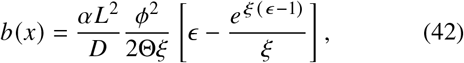

and Eq. (36) allows us to neglect the *ϵ* in the exponent, at which point Eq. (37) allows us to neglect the exponential term altogether, giving

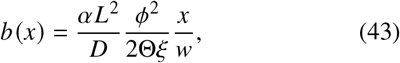

where we have once again restored *ϵ* = *x w*. Eq. (43) is the same as Eq. (40), showing that the bulk profile is insensitive to the choice of boundary condition.

## APPENDIX C

Here we derive Eqs. (18) and (19). Eq. (16) in steady state is solved by directly integrating to find *∂*_*x*_ *b* and integrating again to find *b (x)*. The boundary conditions in Eq. (17) set the integration constants, giving

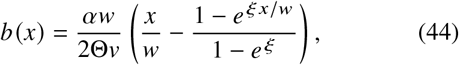

where *ξ* = *vw /D*. With *ϕ* = *w/ L*, Eq. (44) becomes Eq. (18).

As in Appendix A, we write the steady state solution to Eq. (13) as the sum of a particular solution and the homogeneous solution. For the particular solution we use Eq. (4). For the homogeneous solution *c*_0_ = *X (x) Y (y)*, we separate variables as in Appendix A, but here using sinusoidal functions in *x* and exponential functions in *y* is more conducive to the boundary conditions. Specifically,

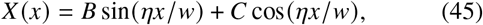

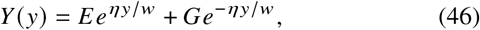

for a constant *η* that arises from the separation. The boundary conditions in Eq. (12) require that *B* = 0 and *η* = *nπ* for nonnegative integer *n*, respectively. Symmetry in *y* requires that *E* = *G*. Thus, the steady state solution to Eq. (13) reduces to

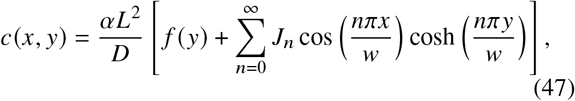

for some unknown *J*_*n*_, where *f(y)* is as in Eq. (4). The boundary condition in Eq. (15) requires

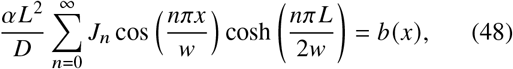

where *b* (*x*) is as in Eq. (44), and we have used the fact that *f* (±*L*/2) = 0. The orthogonality of cosines, expressed as

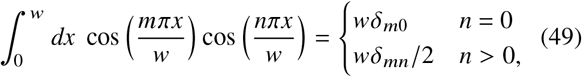

allows us to invert Eq. (48),

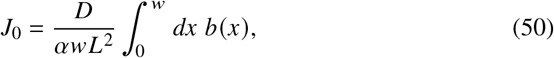

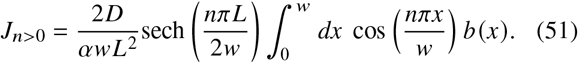

Inserting *b (x)* from Eq. (44) and evaluating the integrals yields

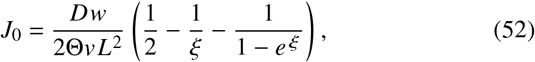

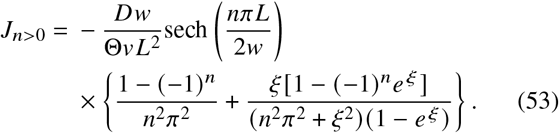

With *ξ* = *vw /D* and *ϕ* = *w /L*, Eqs. (47), (52), and (53) become Eqs. (19), (20), and (21), respectively.

## AUTHOR CONTRIBUTIONS

J. J. W., M. R. B., W. O., K. J., and A. M. designed the research, J. J. W., S. S., and J. E. performed the research and contributed analytic tools, and J. J. W., S. S., W. O., K. J., and A. M. wrote the manuscript.

## ACKNOWLEDGMENTS

This work was supported by National Science Foundation grants DMS-1816315 (W. O.), MCB-1936774 (M. R. B.), MCB-1936770 (K. J.), and MCB-1936761 (A. M.), and by National Institutes of Health grant R01-GM144959 (M. R. B. and K. J.).

## DECLARATION OF INTERESTS

The authors declare no competing interests.

## Notes

### Competing Interest Statement

The authors have declared no competing interest.

